# Harnessing glucocorticoid receptor antagonization to enhance the efficacy of cardiac regenerative growth factors and cytokines

**DOI:** 10.1101/2025.01.15.632969

**Authors:** Silvia Da Pra, Stefano Boriati, Carmen Miano, Irene Del Bono, Francesca Sacchi, Chiara Bongiovanni, Alla Aharonov, Nicola Pianca, Riccardo Tassinari, Carlo Ventura, Mattia Lauriola, Eldad Tzahor, Gabriele D’Uva

## Abstract

Severe myocardial injuries in mammals lead to cardiomyocyte loss and heart failure. Activation of Glucocorticoid Receptor (GR) by endogenous glucocorticoids limit cardiomyocyte proliferation and cardiac regeneration. Here, we reveal that glucocorticoids suppress the proliferative response of cardiomyocytes to a diverse range of regenerative factors, including Neuregulin 1 (NRG1), FGF1, IGF2, IGF1, BMP7, Oncostatin M (OSM), LIF, RANKL, IL6, IL13, IL4, and IL1-beta.

Transcriptomic analyses revealed that glucocorticoids-GR axis induces the expression of MAPK/ERK pathway inhibitors ERRFI1/MIG6 and DUSP1. Using NRG1 as a model system for cardiac regenerative factors, we demonstrate that glucocorticoids suppress growth factor-induced ERK activation, nuclear translocation, and transcriptional output. Knockdown experiments confirmed that ERRFI1 and DUSP1 are key mediators of glucocorticoid-mediated suppression of cardiomyocyte proliferation induced by growth factors. The cardiac expression of GR target genes, including *Dusp1* and *Errfi1,* increases during early postnatal development, coinciding with the transition to post-mitotic cardiomyocytes. In juvenile and adult stage, regenerative growth factors failed to activate MAPK/ERK pathway and induce proliferation effectively; however, DUSP1 inhibition rescues these responses. Consistently, GR antagonization during early postnatal development preserved MAPK/ERK pathway activity. Importantly, GR antagonization or deletion reinstated growth-factor-induced proliferation of postmitotic cardiomyocytes. Finally, GR antagonization enhanced the *in vivo* effectiveness of growth factor-based regenerative therapy in an adult mouse model of myocardial injury, enhancing cardiomyocyte proliferation and cardiac function. These findings reveal a previously unrecognized mechanism by which systemic hormones, specifically glucocorticoids, regulate the regenerative potential of cardiac paracrine factors. This study highlights transient glucocorticoid inhibition as a promising strategy to enhance the therapeutic efficacy of growth factor-based cardiac regenerative approaches.

**Highlights:**

- Physiological glucocorticoids suppress the proliferative response of cardiomyocytes to regenerative growth factors and cytokines.
- Glucocorticoids inhibit growth factor-induced MAPK/ERK activation in cardiomyocytes by upregulating the expression of negative regulators DUSP1 and ERRFI1.
- Increased glucocorticoid-induced expression of DUSP1 and ERRFI1 during early postnatal development impairs the mitogenic activity of cardiac regenerative factors.
- GR antagonization restores growth factor-induced MAPK signaling and mitogenic activity in postmitotic cardiomyocytes and enhances the efficacy of growth factor-based regenerative therapies following myocardial injury.

## Introduction

Nowadays, cardiovascular diseases are the leading cause of death worldwide (Martin et al., 2024). Despite their heterogeneity in causes, manifestations, severity, and treatment strategies, most myocardial pathologies share the death and permanent loss of cardiac muscle cells (cardiomyocytes) as a common hallmark (Secco & Giacca, 2023) (Martin et al., 2024) (Whelan et al., 2010) (Chiong et al., 2011). Cardiomyocyte loss can arise from acute events, such as myocardial infarction, viral myocarditis (Kytö et al., 2004), or cardiotoxic side effects of anticancer treatments like anthracyclines (Y.-W. Zhang et al., 2009) (Octavia et al., 2012), or chronic conditions, including hypertensive heart disease (Gonzalez, 2003), aortic stenosis (Hein et al., 2003), arrhythmogenic cardiomyopathy (Yamaji et al., 2005) and inherited cardiac disorders (Hashem et al., 2015). Given the limited regenerative capacity of the adult mammalian heart, cardiomyocytes loss significantly contributes to cardiac dysfunction and heart failure (Bongiovanni et al., 2021) (Tzahor & Poss, 2017) (Sadek & Olson, 2020) (Eschenhagen et al., 2017), presenting a major clinical challenge.

Enhancing the proliferative capacity of endogenous cardiomyocytes holds promise for regenerating heart tissue and restoring cardiac function (Tzahor & Poss, 2017) (Sadek & Olson, 2020) (van Berlo & Molkentin, 2014) (Hashimoto et al., 2018) (Uygur & Lee, 2016). Among proposed strategies, the administration of specific growth factors and cytokines has shown potential to stimulate cardiomyocyte proliferation and promote heart regeneration. For instance, administration of Neuregulin 1 has been suggested to induce cardiomyocyte proliferation and heart regeneration in adult mice (D’Uva et al., 2015) (Bersell et al., 2009). However, the mitogenic activity of NRG1 declines markedly during early postnatal development (D’Uva et al., 2015) (Polizzotti et al., 2015). This effect is primarily consequent to reduced expression of its coreceptor ERBB2 (D’Uva et al., 2015). Indeed, overexpression of a constitutively active ERBB2 (caERBB2) isoform strongly induces cardiomyocyte dedifferentiation and proliferation, even at juvenile or adult non-regenerative stages (D’Uva et al., 2015). In line with the limited mitogenic activity on cardiomyocyte at adult stage, clinical trials testing NRG1 administration for chronic heart failure have shown modest but yet favourable effects on cardiac function (Gao et al., 2010) (Jabbour et al., 2011) (Lenihan et al., 2016). Additional regenerative growth factors identified in preclinical studies include FGF1 (Engel et al., 2006) (Novoyatleva et al., 2014) (Engel et al., 2005), IGF2 (Shen et al., 2020) (Schuetz et al., 2024b), IGF1 (Sundgren et al., 2003a) (Schuetz et al., 2024b) (Koudstaal et al., 2014), BMP7 (Bongiovanni, Bueno-Levy, et al., 2024), Oncostatin M (OSM) (Kubin et al., 2011) (Y. Li et al., 2020), LIF (Kubin et al., 2011) (Zou et al., 2003) (Negoro et al., 2001), RANKL (Zacchigna et al., 2018) (Ock et al., 2012), IL6 (Tang et al., 2018) (C. Han et al., 2015a) (Zogbi et al., 2020) (Fahmi et al., 2013), IL13 (Wodsedalek et al., 2019) (Paddock et al., 2021), IL4 (Paddock et al., 2021) (Zogbi et al., 2020), IL1-β (Palmer et al., 1995) (C. Han et al., 2015a) (Engel et al., 2005) (Przybyt et al., 2013) (Singh et al., 2016). Recently, we highlighted glucocorticoids as key hormones driving cardiomyocyte maturation and cell cycle exit during early postnatal development, thereby contributing to the decline in regenerative potential observed in juvenile and adult stages (Pianca et al., 2022). Although the effects of glucocorticoids and growth factors have been studied separately in cardiac regenerative medicine, others and we have previously shown that their molecular interplay is crucial in the regulation of cellular outcomes in other contexts (D’Uva & Lauriola, 2016) (Lauriola et al., 2014).

In this study, we investigated the impact of physiological glucocorticoids on cardiomyocyte proliferation and regeneration driven by cardiac regenerative factors. Our findings propose a model where glucocorticoids dampen MAPK/ERK activation in cardiomyocytes by modulating signal intensity, localization, and transcription of downstream effectors. This inhibition significantly curbs the potential of multiple cardiac regenerative growth factors to stimulate cardiomyocyte proliferation. We suggest transient pharmacological antagonization of glucocorticoid receptor activity to enhance the regenerative efficacy of growth factors, facilitating cardiomyocyte replacement thus improving recovery after cardiac injuries.

## Results

### Glucocorticoids suppress the mitogenic effects of regenerative growth factors on cardiomyocytes

To test the impact of physiological glucocorticoids on cardiomyocyte proliferation and regeneration driven by cardiac regenerative factors, we assessed the proliferation of cultured neonatal murine cardiomyocytes following *in vitro* administration of several cardiac regenerative growth factors, including NRG1, FGF1, IGF2, IGF1, BMP7, OSM, LIF, RANKL, IL6, IL13, IL4, and IL1-beta, in combination with corticosterone, the primary glucocorticoid in rodents. Cardiomyocyte proliferation was evaluated by BrdU incorporation coupled with immunostaining for a cardiomyocyte-specific marker Troponin I (cTnI). Remarkably, our findings demonstrated that corticosterone treatment suppresses the mitogenic effects of all the regenerative growth factors evaluated (**Fig. 1a**). To deepen our understanding, we used NRG1 as the major model system. Analysis of Aurora B Kinase immunoreactivity, a marker of midbody formation during cytokinesis, showed that corticosterone treatment suppresses NRG1-induced cardiomyocyte cytokinesis (**Fig. 1b**). The inhibitory effect of corticosterone on NRG1-induced cardiomyocyte division was further confirmed using time-lapse imaging of cardiomyocytes identified by TMRE (tetramethylrhodamine ethyl ester) staining (Hattori et al., 2010) (**Fig. 1c**).

**Figure 1.**
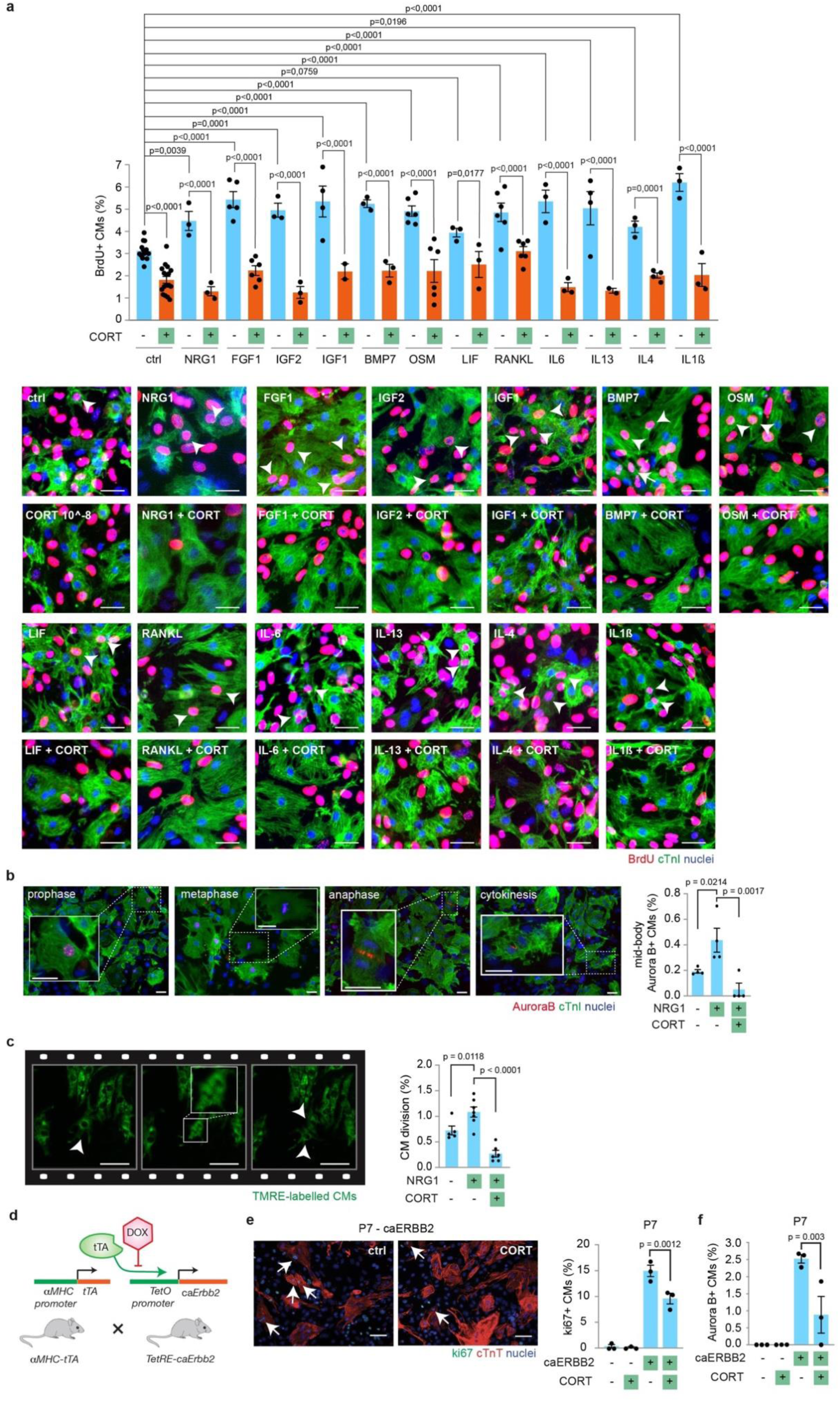
Glucocorticoids suppress the mitogenic activity of cardiac regenerative growth factors and cytokines. (**a**) Immunofluorescence analysis of DNA synthesis of post-natal day 1 (P1) cardiomyocytes cultured *in vitro* and stimulated for 48h with selected growth factors, namely NRG1 (100 ng/mL), FGF1 (10 ng/mL), IGF2 (10 ng/mL), IGF1 (10 ng/mL), BMP7 (10 ng/mL), OSM (10 ng/mL), LIF (10 ng/mL), RANKL (10 ng/mL), IL6 (10 ng/mL), IL13 (10 ng/mL), IL4 (10 ng/mL), IL1-beta (10 ng/mL), alone or in combination with corticosterone (CORT, 10^−8^ M) (n = 40984 cardiomyocytes pooled from the analysis of 133 samples). Cardiomyocytes were identified by cardiac Troponin I (cTnI) staining and analysed by immunofluorescence for DNA synthesis (BrdU incorporation assay). Representative figures are provided; white arrows point to cardiomyocytes undergoing DNA synthesis; scale bar 50 μm; (**b**) Immunofluorescence analysis of cytokinesis of post-natal day 1 cardiomyocytes cultured *in vitro* and stimulated for 48h with corticosterone (CORT, 10-8 M), alone or in combinations with NRG1 (100ng/ml) (n = 7833 cardiomyocytes pooled from the analysis of 12 samples). Cardiomyocytes were identified by cardiac Troponin I (cTnI) staining and analyzed by immunofluorescence for cytokinesis (midbody Aurora B kinase). Representative figures are provided; white arrows point to Aurora B positive cardiomyocytes; scale bar 30 μm; (**c**) Quantification and representative images of cell division events in TMRE-labeled neonatal cardiomyocytes detected in 16-h time-lapse imaging at 20-min intervals *in vitro* (n = 18 samples with a total of 14239 cardiomyocytes analyzed). Representative figures are provided; arrows point at cardiomyocytes undergoing cell division; scale bars, 50 μm; **(d)** Diagram depicting the cross-breeding to generate a constitutively active ERBB2 overexpression (caERBB2) in cardiomyocytes; **(e-f**) Immunofluorescence analysis of (**e**) cell-cycle re-entry (Ki67) and (**f**) cytokinesis (Aurora B kinase) of control and caERBB2 post-natal day 7 (P7) cardiomyocytes following stimulation with corticosterone (CORT, (n = 1667 cardiomyocytes pooled from the analysis of 12 samples for **e**; n = 1850 cardiomyocytes pooled from the analysis of 12 samples for **f**). Representative images are provided; arrows point at cycling cardiomyocytes; scale bars,60 μm). The values are presented as mean (error bars show SEM), statistical significance was determined using one-way ANOVA followed by Sidak’s test in (**a**), (**b**), (**c**), (**e**) and (**f**) (comparison between pairs of treatments).

The mitogenic activity of NRG1 declines markedly during early postnatal development (D’Uva et al., 2015) (Polizzotti et al., 2015), primarily due to a reduction in the expression of its coreceptor ERBB2 (D’Uva et al., 2015). Notably, overexpression of a constitutively active ERBB2 (caERBB2) isoform strongly promotes cardiomyocyte dedifferentiation and proliferation, even in juvenile or adult stages when the heart has lost its intrinsic regenerative capacity (D’Uva et al., 2015). Using a transgenic mouse model with cardiomyocyte-specific induction of a constitutively active ERBB2 (ca*ERBB2* mice, **Fig. 1d**) (D’Uva et al., 2015), we investigated the impact of glucocorticoids on ERBB2-induced proliferative capacity in juvenile, postmitotic cardiomyocytes. Remarkably, treatment of cultured cardiomyocytes isolated from ca*ERBB2* mice at postnatal day 7 with corticosterone significantly reduced ERBB2-induced cardiomyocyte cell cycle re-entry (**Fig. 1e**) and division (**Fig. 1f**). In summary, our screening reveals that glucocorticoids inhibit cardiomyocyte proliferation stimulated by a wide array of cardiac regenerative growth factors, cytokines, and potent downstream mitogenic mediators.

### GR activation induces the expression of negative regulators of the MAPK/ERK signalling in cardiomyocytes

To gain mechanistic insights into the cytostatic effect induced by glucocorticoids on cardiac regenerative growth factor signalling, we performed whole transcriptome RNA sequencing on primary neonatal cardiomyocytes treated *in vitro* with corticosterone. Corticosterone was administrated over a short time frame to identify genes directly regulated by glucocorticoids. Our analysis identified a significant upregulation (adjusted p value < 0.05) of 40 genes following corticosterone treatment (full list provided in **Supplementary table 1**). Gene Ontology analysis of transcriptomic data revealed an enrichment of genes involved in the “Negative regulation of the MAPK cascade” (**Fig. 2a**), a pathway previously shown to mediate the proliferative and regenerative activity of the tested growth factors (Bersell et al., 2009) (D’Uva et al., 2015) (Engel et al., 2006) (Novoyatleva et al., 2014) (Engel et al., 2005) (Shen et al., 2020) (Schuetz et al., 2024b) (Sundgren et al., 2003a) (Koudstaal et al., 2014) (Bongiovanni, Bueno-Levy, et al., 2024) (Kubin et al., 2011) (Y. Li et al., 2020) (Zou et al., 2003) (Negoro et al., 2001) (Zacchigna et al., 2018) (Ock et al., 2012) (Tang et al., 2018) (C. Han et al., 2015a) (Zogbi et al., 2020) (Fahmi et al., 2013) (Wodsedalek et al., 2019) (Paddock et al., 2021) (Palmer et al., 1995) (Przybyt et al., 2013) (Singh et al., 2016). The enriched genes in the Gene Ontology term “Negative regulation of MAPK cascade”, namely ERRFI1, DUSP1, RGS2, and PER1, were also identified among the top 20 most significantly upregulated genes in response to corticosterone stimulation (**Fig. 2b**). Notably, ERRFI1 and DUSP1 are well-recognized direct inhibitors of key MAPK components. ERRFI1, also known as MIG-6 or RALT, is a cytosolic protein inhibiting the catalytic activation of ERBB receptors, and promoting their lysosomal degradation (Citri & Yarden, 2006) (Segatto et al., 2011). DUSP1, a member of the dual-specificity phosphatases (DUSPs) family, is a nuclear enzyme that inactivates all three branches of the MAPK axis, namely ERK, JNK and p38 (Keyse, 2008). Importantly, glucocorticoid-induced expression of ERRFI1 and DUSP1 has been shown to suppress MAPK/ERK signalling in response to the growth factor EGF in quasi-normal breast cells (Lauriola et al., 2014). To further characterize the kinetics of *Dusp1* and *Errfi1* induction in cardiomyocytes, we measured their mRNA expression levels over time following corticosterone treatment. *Dusp1* expression increased rapidly, peaking at 1.5 hours post-treatment **(Fig. 2c**), while *Errfi1* induction markedly increased at 2 hours and peaked at 4 hours post-treatment **(Fig. 2d**). At protein level, we confirmed that corticosterone activates GR **(Fig. 2e).** Notably, the prolonged stimulation for 12-24 hours led to a decrease in GR abundance **(Fig. 2e**), consistent with previous studies showing reduced stability of activated GR over time (Wallace & Cidlowski, 2001). DUSP1 protein, which exhibited low basal expression in neonatal cardiomyocytes, was significantly upregulated at 6 and 12 hours following corticosterone stimulation, with levels declining thereafter **(Fig. 2e**). This transient increase aligns with earlier studies suggesting that DUSP proteins undergo rapid turnover (Kassel et al., 2001) (Chen et al., 2019). In contrast, following corticosterone treatment, ERRFI1 levels exhibited a modest increase as early as 2 hours and remained stable up to 24 hours **(Fig. 2e**). Pre-treatment with GR antagonist, RU486 (Mifepristone), completely abolished the corticosterone*-*induced upregulation of *Dusp1* and *Errfi1* expression **(Fig. 2f-g**), confirming GR as the mediator of this effect. To further evaluate the role of GR, we analysed *Dusp1* and *Errfi1* expression in a cardiomyocyte-specific knock-out mouse model generated by Cre-Lox technology (**Fig. 2h**) (Pianca et al., 2022). Neonatal cardiomyocytes isolated from GR-ablated and control mice were cultured *in vitro* and stimulated with corticosterone. In GR-deficient cardiomyocytes, corticosterone-induced expression of *Errfi1* and *Dusp1* mRNA was decreased, confirming that GR is essential for the upregulation of these genes **(Fig 2i-j**). Collectively, these results demonstrate that GR activation by glucocorticoids in cardiomyocytes drives the expression of Dusp1 and Errfi1, key negative regulators of the MAPK/ERK pathway.

**Figure 2.**
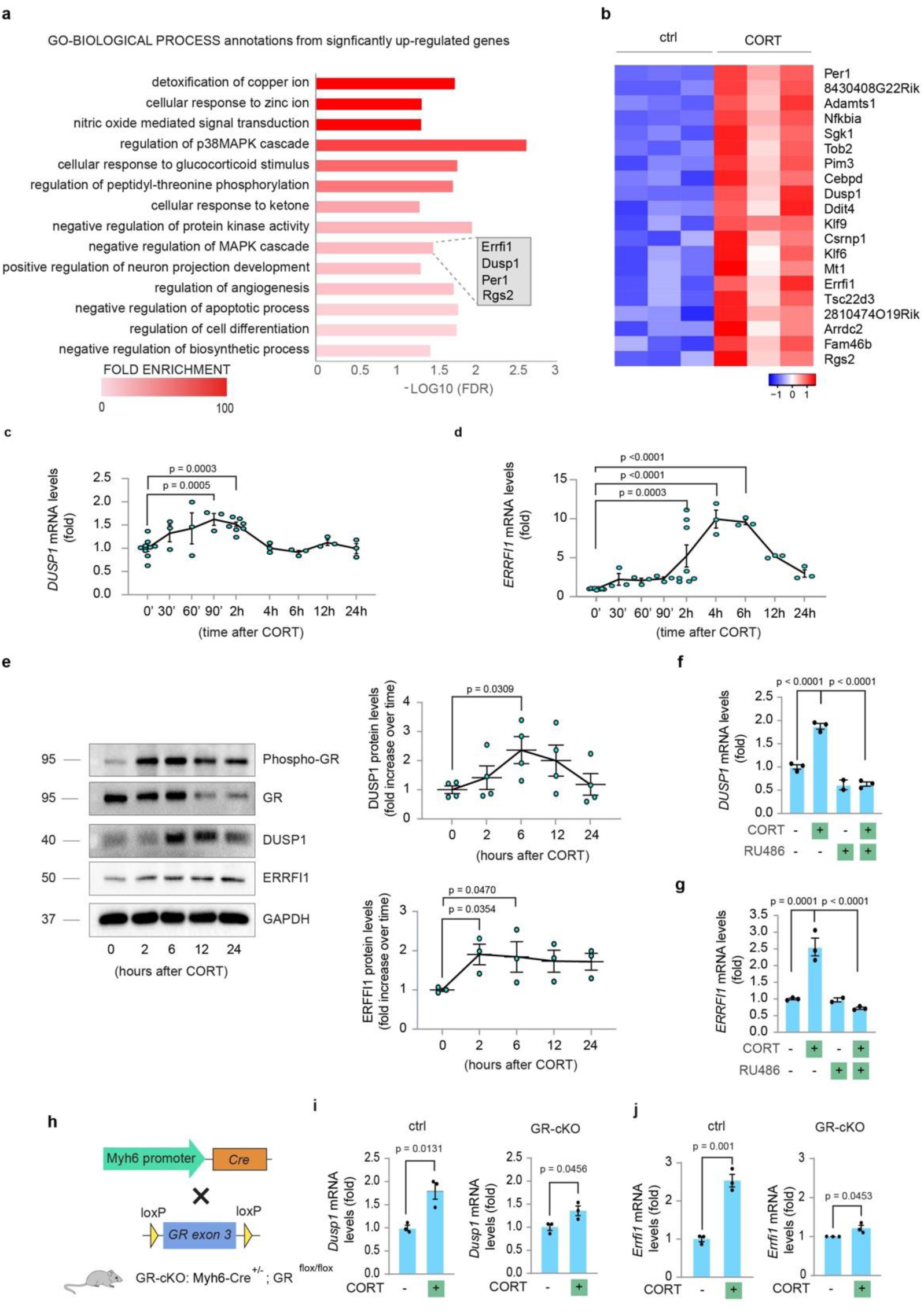
Glucocorticoids trigger the expression of negative regulators of MAPK signalling pathway via GR activation. (**a**) Gene Ontology analysis of transcriptomic data from cultured neonatal cardiomyocytes, separated from stromal cells by immunomagnetic separation, upon *in vitro* treatment with corticosterone (CORT, 10^−6^ M) for 30 minutes. The box shows induced genes annotated within the GO term “negative of MAPK cascade” (GO:0043409); (**b**) Heatmap of the top 20 most significantly up-regulated genes (Adj pvalue < 0.05, LogFC > 0) by RNA-Seq analysis of cultured neonatal enriched cardiomyocytes, separated from stromal cells by immunomagnetic separation, treated *in vitro* with corticosterone (CORT, 10^−6^ M) for 30 minutes (n = 3 replicates per condition); **(c, d)** RT-qPCR analysis of mRNA expression levels of *Errfi1* and *Dusp1* in neonatal enriched cardiomyocyte, separated from stromal cells by immunomagnetic separation, upon *in vitro* treatment with corticosterone (CORT, 10^−6^ M) at different timepoints (0’, 30’, 60’, 90’, 2H, 4H, 6H, 12H, 24H) (n = 9 replicates) for 0’, 3 replicates for 30’, 60’, 90’, 6H, 12H, 24H and 7 replicates for 2H); **(e)** Western blot analysis of GR, phospho-GR, DUSP1 (n = 4 replicates per timepoint) and ERRFI1 (n = 3 replicates per timepoint) protein levels upon stimulation with corticosterone (CORT, 10^−6^ M) at different timepoints (2H, 6H, 12H and 24H) in neonatal enriched cardiomyocytes, separated from stromal cells by immunomagnetic separation; **(f-g)** RT-qPCR analysis of mRNA expression of **(f)** *Dusp1* and **(g)** *Errfi1* upon pre-treatment for 30 min with the GR-antagonist RU486 (10^−5^ M) and/or treatment with corticosterone (CORT, 10^−6^ M) for 2 additional hours in neonatal enriched cardiomyocytes, separated from stromal cells by immunomagnetic separation; **(h**) Schematic diagram depicting the cross-breeding to generate a GR-cKO mice; (**i-j**) RT-qPCR analysis of mRNA expression levels of **(i**) *Dusp1* and (**j**) *Errfi1* in neonatal cardiomyocyte cultures isolated from control (ctrl) and cardiomyocyte-restricted GR knock-out (GR-cKO) mouse model upon *in vitro* treatment with corticosterone (CORT, 10^−6^ M) for 30 minutes (n = 3 replicates per condition). The values are presented as mean (error bars show SEM), statistical significance was determined using one-way ANOVA followed by Sidak’s test in **(c)**, **(d)**, **(e**), (**f**) and **(g)** (comparison between pairs of treatments), and two-sided Student’s t-test in (**i)** and (**j**).

### Glucocorticoids restrain growth-factor-induced MAPK/ERK activation, nuclear localization and transcriptional outcome in cardiomyocytes

Using NRG1 as a model system of cardiac regenerative growth factors acting through the ERK/MAPK pathway, we investigated whether glucocorticoids and their induced genes DUSP1 and ERRFI1 contribute to the suppression of growth-factor-induced ERK signalling in cardiomyocytes.

To begin, we confirmed that corticosterone-induced genes DUSP1 and ERRFI1 indeed repress ERK signalling in cardiomyocytes. To this end, knock down of *Errfi1* and *Dusp1* was achieved by specific small interfering RNAs (siRNAs), whose efficacy was validated 48 hours post-transfection (**Supplementary Figs. 1a, b**). Consistent with their role as ERK inhibitors, Dusp1 or Errfi1 knockdown significantly increased ERK1/2 phosphorylation, without affecting AKT phosphorylation **(Fig. 3a**). Next, we analysed ERK activation following NRG1 treatment in primary neonatal cardiomyocytes pre-treated with corticosterone. As hypothesized, corticosterone attenuated the amplitude of NRG1-induced ERK activation **(Fig. 3b**). We hypothesized that corticosterone-induced DUSP1, a nuclear phosphatase, specifically attenuates nuclear ERK activation, which is known to be associated with cell proliferation (Plotnikov et al., 2015) (Michailovici et al., 2014) (Avraham & Yarden, 2011). To test this, we examined the subcellular localization of activated ERK in cardiomyocytes using immunofluorescence for phosphorylated ERK1/2 and a cardiomyocyte-specific marker. Notably, corticosterone pre-treatment abolished the increase in nuclear ERK activation (phosphorylation) induced by NRG1 in cardiomyocytes **(Fig. 3c**). Activation of the MAPK/ERK signalling pathway triggers rapid and transient induction of Immediate Early Genes (IEGs), such as *Fos, Jun-b and Egr1* (Avraham & Yarden, 2011) (Amit et al., 2007). Notably, these genes serve as key regulators of cardiac regenerative capacity. Indeed, EGR1 is essential for heart regeneration in neonatal mice (L. Zhang et al., 2024), and AP-1, a transcription factor family comprising multiple Jun and Fos members, is required for heart regeneration in zebrafish (Beisaw et al., 2020). Consistently, we found that NRG1 transiently upregulates *Fos, Jun-b and Egr1* expression within 30-60 minutes in primary cardiomyocytes **(Supplementary Figs. 2a-c**). Similarly, bioinformatic analysis of RNA*-*sequencing data from cardiac tissue isolated from transgenic mouse model with cardiomyocyte-specific induction of a constitutively active ERBB2 isoform (ca*ERBB2* mice) revealed significantly elevated expression of immediate early ERK target genes, *Egr1, Fos* and *Junb* compared to controls **(Supplementary Figs. 3a-c**). In line with its inhibitory effects on ERK activation and nuclear translocation, corticosterone pre-treatment reduced the induction of *Fos, Junb and Egr1* in response to NRG1 stimulation (**Fig. 3d**). Together, these results demonstrate that activation of glucocorticoid receptor by physiological glucocorticoids reduces growth factor-induced ERK activation, deactivates nuclear ERK and inhibits the transcription of its target genes in cardiomyocytes.

**Figure 3.**
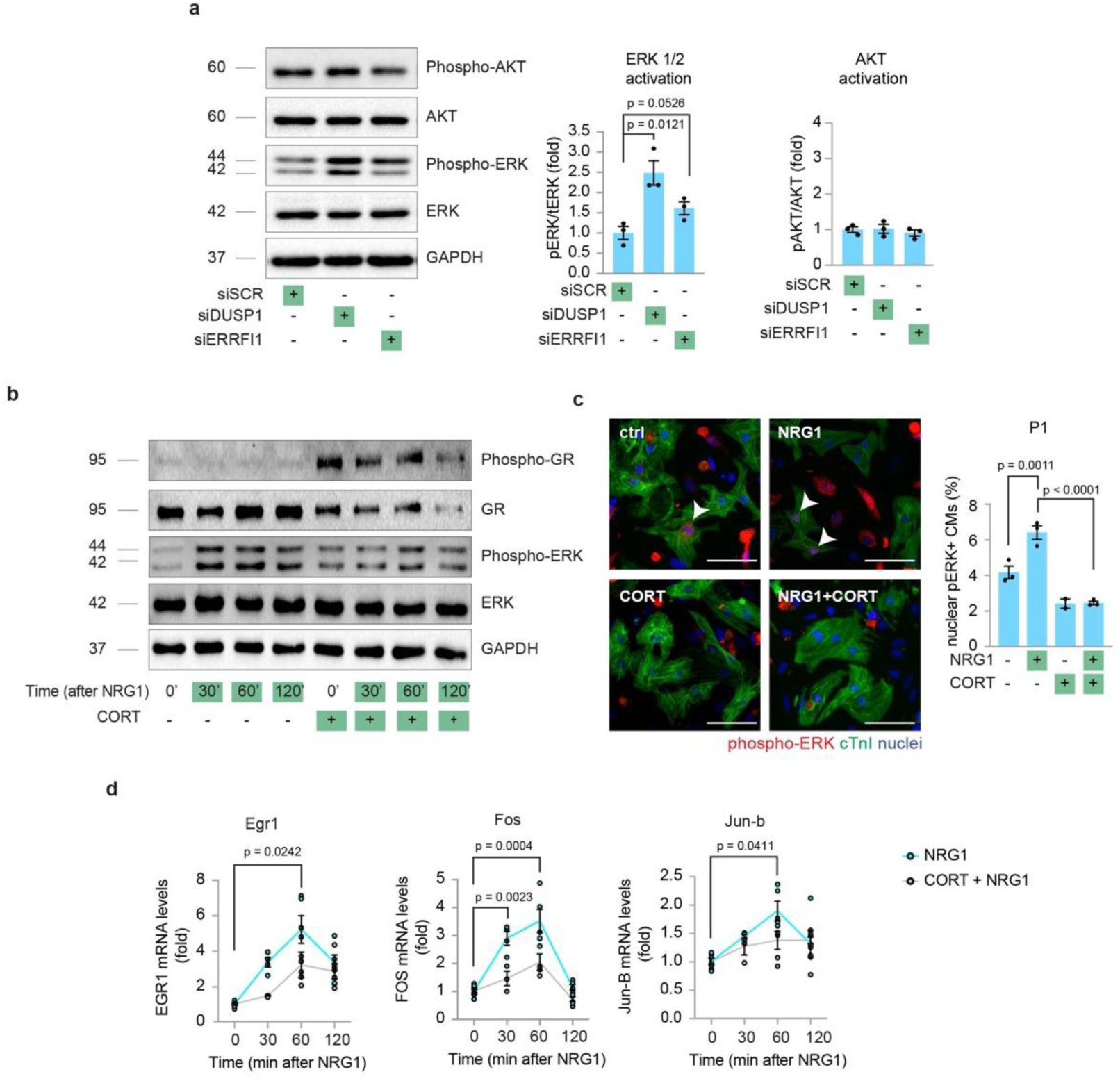
Glucocorticoids intercept growth-factor-induced MAPK/ERK signalling activation in cardiomyocytes. (**a)** Western Blot analysis and quantification of AKT and ERK activation (phosphorylation) upon silencing of *Dusp1* or *Errfi1* mRNA expression through transfection of a specific siRNA in neonatal enriched cardiomyocytes, separated from stromal cells by immunomagnetic separation (n = 3 replicates per condition); GAPDH protein levels are provided as second loading control; (**b**) Analysis of GR and ERK activation by western blot analysis of phospho-GR, total GR, phospho-ERK 1/2, total ERK and GAPDH protein levels, in enriched neonatal cardiomyocytes, separated from stromal cells by immunomagnetic separation, following stimulation with NRG1 (100 ng/ml) for 30, 60 and 120 minutes with/without 6 hours pre-treatment with corticosterone (CORT, 10^−6^ M) (n = 2 replicates per condition); (**c**) Immunofluorescence analysis of phospho-ERK nuclear translocation upon treatment with NRG1 (100ng/ml) for 15 min with/without 6 hours pre-treatment with corticosterone (CORT, 10^−6^ M) in mixed culture of neonatal cardiomyocytes (n = 4610 cardiomyocytes pooled from the analysis of 11 samples). Cardiomyocytes were identified by cardiac Troponin I (cTnI) staining and analyzed by immunofluorescence for pERK nuclear localization. Representative pictures are provided; arrows point at cardiomyocytes with nuclear phospho-ERK immunoreactivity; scale bar 50 μm; (**d**) Real-time PCR analysis of the expression of IEGs (Immediate Early Genes) such as *Fos* (n = 19 replicates for NRG1, 16 replicates for CORT+NRG1)*, Jun-b* (n = 19 replicates for NRG1, 16 replicates for CORT+NRG1), and *Egr1* (n = 19 replicates for NRG1, 17 replicates for CORT + NRG1) upon treatment with NRG1 (100 ng/ml) for 30, 60 and 120 minutes with/without 6 hours pre-treatment with Corticosterone (CORT, 10^−6^ M) in neonatal enriched cardiomyocytes, separated from stromal cells by immunomagnetic separation. The values are presented as mean (error bars show SEM), statistical significance was determined using two-sided Student’s t test in (**a**), using one-way ANOVA followed by Sidak’s test (**c**) and using two-way ANOVA followed by Sidak’s test in **(d)** (comparison between pairs of treatments).

### Increased glucocorticoid-induced DUSP1 and ERRFI1 expression during early postnatal development suppresses the mitogenic activity of cardiac regenerative factors

To assess whether DUSP1 and ERRFI1 mediate the corticosterone-induced suppression of the mitogenic activity of cardiac regenerative factors, we knocked down their expression in neonatal cardiomyocytes using siRNA for 48 hours. After 48 hours of siRNA treatment, cells were exposed to a combination of corticosterone and NRG1 for an additional 48 hours. Silencing *Dusp1* and/or *Errfi1,* either individually or in combination, significantly increased cardiomyocyte proliferation (**Fig. 4a**), highlighting their role as brakes on cardiomyocyte proliferation. Importantly, corticosterone’s suppression of NRG1-induced cardiomyocyte proliferation was completely abolished when *Dusp1* and *Errfi1* were silenced (**Fig. 4a).** These data demonstrate that glucocorticoids inhibit NRG1-induced cardiomyocytes proliferation through *Dusp1* or *Errfi1*. Unexpectedly, in the absence of NRG1, silencing *Dusp1* or *Errfi1* did not prevent the corticosterone’s suppressive effect on cardiomyocyte proliferation (**Fig. 4a**). Thus, the suppression of cardiomyocyte proliferation by glucocorticoids in the absence of growth factor stimulation appears to be independent of DUSP1 and ERRFI1, potentially involving other yet-to-be-identified molecular players. In brief, these data suggest that DUSP1 and ERRFI1 specifically mediate the glucocorticoid-induced suppression of mitogenic activity triggered by growth factor-induced MAPK/ERK activation.

**Figure 4.**
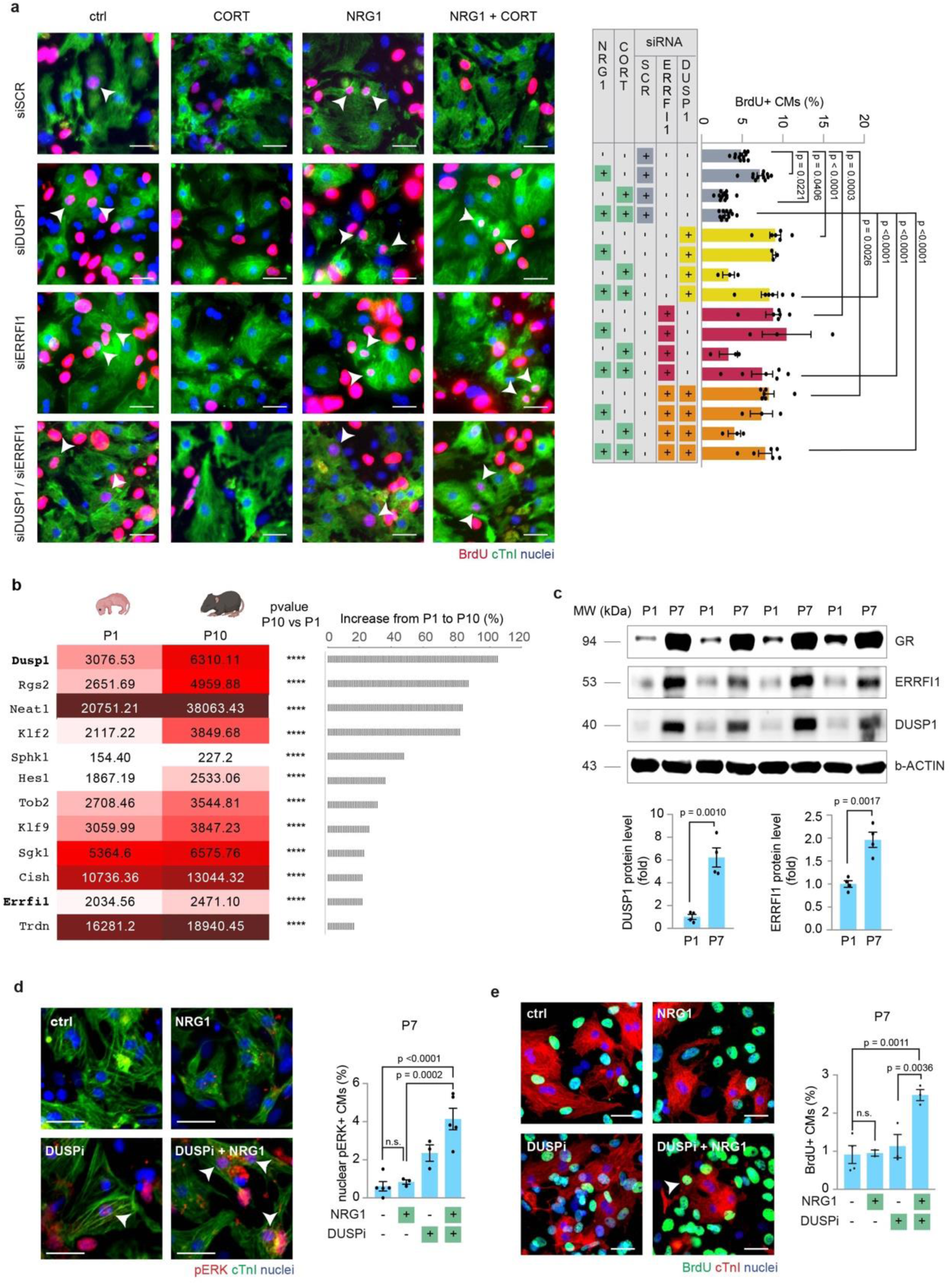
Increased glucocorticoid-induced DUSP1 and ERRFI1 expression in early postnatal development progressively impairs the mitogenic potential of regenerative factors. **(a)** Immunofluorescence analysis of DNA synthesis (BrdU incorporation,) of mixed cultures of neonatal cardiomyocytes following knock-down of Dusp1 and/or Errfi1 with/without stimulation with NRG1 (100ng/ml) and/or CORT (10^−8^ M). Cardiomyocytes were identified by cTnI staining and analysed by immunofluorescence for DNA synthesis (BrdU) (n = 33676 cardiomyocytes pooled from the analysis of 102 samples); **(b)** mRNA expression levels of GR targets in mouse postnatal day 1 (P1, first column) and post-natal day 10 (P10, second column) heart lysates obtained by meta-analysis of RNA-seq data (Haubner et al, 2012), along with the calculated increase from P1 to P10 in terms of percentage (graph bar). The values in the first and second columns are presented as mean expression levels of three biological replicates; the p value is reported for the increase from P1 to P10 (third column) as follows *p % 0.05; **p % 0.01; ***p % 0.001; ****p % 0.0001; **(c)** Western blot analysis of total protein expression of GR, DUSP1 and ERRFI1/MIG6 in mouse postnatal day 1 (P1) and postnatal day 7 (P7) heart lysates (n = 4 mice per timepoint); b-Actin protein levels are provided as second loading control; **(d-e)** Immunofluorescence analysis of DNA synthesis (BrdU) in post-natal day 7 (P7) cultured cardiomyocytes following stimulation with DUSP1-inhibitor (BCI) (10^−7^ M), alone or in combinations with NRG1 (100ng/ml). Cardiomyocytes were identified by cTnI staining and analysed by immunofluorescence for (**d**) pERK nuclear localization (n = 4271 cardiomyocytes pooled from the analysis of 16 samples), or (**e**) DNA synthesis (BrdU) (n = 2827 cardiomyocytes pooled from the analysis of 12 samples). Representative pictures are provided; arrows point at proliferating CMs; scale bars, 30 μm; arrows point at cardiomyocytes with (**d**) nuclear phospho-ERK immunoreactivity or (**e**) undergoing DNA synthesis; scale bars, 30 μm. The values are presented as mean (error bars show SEM), statistical significance was determined using one-way ANOVA followed by Dunnet’s test in (**a**), using two-sided Student’s t test in (**c**) and using one-way ANOVA followed by Sidak’s test in (**d**), (**e**).

The activity of glucocorticoids-GR axis has been shown to increase during the early postnatal period (Pianca et al., 2022). We thus decided to evaluate the role of DUSP1 and ERRFI1 at later postnatal stages. By intersecting GR target genes in cardiomyocytes (**Supplementary Table 1**) with genes increasing their expression during early postnatal cardiac development (Haubner et al., 2012), we here show that several GR-induced genes, including *Dusp1* and *Errfi1,* are upregulated during the early postnatal development (**Fig. 4b**). Western blot analysis revealed a marked increase in DUSP1 and ERRFI1 protein expression during this period (**Fig. 4c**). Using cultured cardiomyocytes isolated at a later developmental stage (P7), we observed that NRG1 failed to induce significant phospho-ERK nuclear translocation or cell proliferation in P7 cardiomyocytes (**Fig. 4d-e**). This contrasts with the robust nuclear ERK activation and proliferative response observed in neonatal cardiomyocytes (**see Fig. 3c and 1a**). These findings are consistent with previously reported reductions in the mitogenic activity of NRG1 at this developmental stage (D’Uva et al., 2015). Remarkably, pharmacological inhibition of DUSP1 with its allosteric inhibitor BCI restored NRG1-induced nuclear ERK activation and cardiomyocyte proliferation in P7 cardiomyocytes (**Fig. 4d-e**). These data suggest that the decline in ERBB2 expression (D’Uva et al., 2015) and the upregulation of DUSP1 expression (this study) both contribute to limiting the mitogenic potential of NRG1 during early postnatal development.

Overall, these findings demonstrate that glucocorticoid-mediated suppression of growth factor-induced mitogenic signaling in cardiomyocytes is driven by the upregulation of the MAPK/ERK inhibitors ERRFI1 and DUSP1. Moreover, they suggest that heightened glucocorticoid activity during the early postnatal period increases the expression of ERRFI1 and DUSP1, thereby suppressing growth factor-induced ERK signaling and the associated mitogenic response.

### GR antagonization in post-mitotic cardiomyocytes restores their responsiveness to growth factor mitogenic stimuli

Based on our data, which show that increased glucocorticoid-mediated expression of DUSP1 and ERRFI1 suppresses growth factor-induced ERK signaling and cell proliferation, we hypothesize that inhibiting the glucocorticoid-GR axis can restore growth factor-induced signaling and mitogenic activity during juvenile and adult stages. Supporting this hypothesis, in vivo GR antagonization from birth to postnatal day 7 was sufficient to sustain ERK signaling in cardiomyocytes, as evidenced by increased nuclear localization of ERK (**Fig. 5a**). Importantly, cardiomyocyte-specific GR deletion or transient GR antagonization using RU486 rescued NRG1-induced cell cycle re-entry and proliferation in cardiomyocytes at the post-mitotic juvenile stage (postnatal day 7 - P7) (**Figs. 5b-c)**. Since cardiomyocytes at postnatal day 7 are not fully differentiated or entirely post-mitotic, we extended our study to include cardiomyocytes derived from adult mice. In this model, NRG1 failed to induce cardiomyocyte proliferation, while combinatorial treatment resulted in a significant increase in cardiomyocyte proliferation, further supporting the synergistic efficacy of this strategy (**Fig. 5d**).

**Figure 5.**
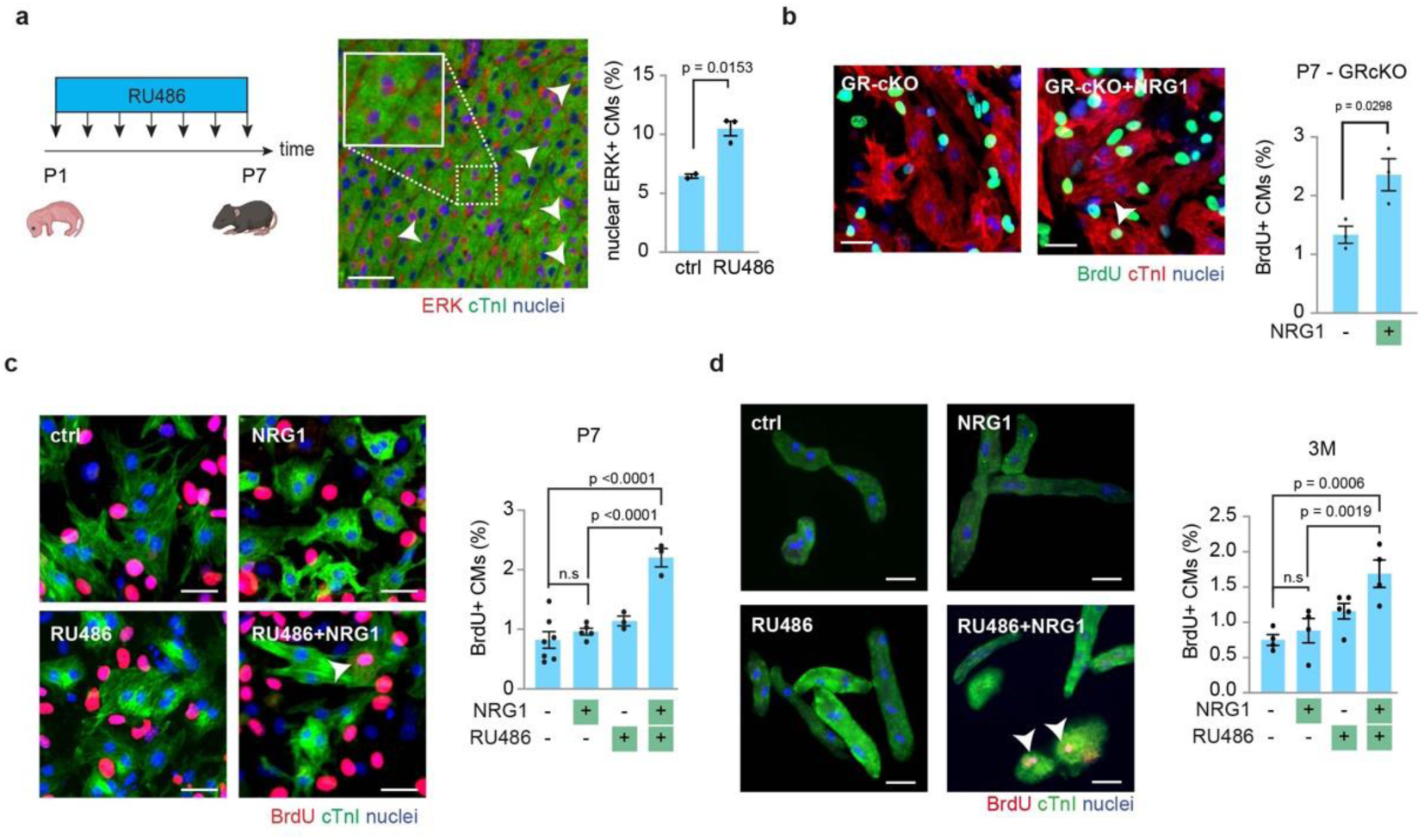
Inhibition of the GCs/GR axis rescues the mitogenic potential of regenerative factors in post-mitotic cardiomyocytes. **(a)** In vivo evaluation of MAPK activity by quantification of nuclear localization of total ERK and cTnI staining in heart sections obtained from P7 mice with and without administration of RU486 in the drinking water starting from birth. Cardiomyocytes were identified by cTnI staining and analysed by immunofluorescence for nuclear ERK localization (n = 5 mice with a total 4488 CMs analyzed). Representative images are provided. Arrows point at CMs positive for nuclear ERK; scale bars, 20 μm; **(b)** Immunofluorescence analysis of DNA synthesis (BrdU) of GR-cKO post-natal day 7 (P7) cardiomyocytes following stimulation with NRG1 (100ng/ml) (n = 1236 cardiomyocytes pooled from the analysis of 6 samples). Cardiomyocytes were identified by cTnI staining and analysed by immunofluorescence for DNA synthesis (BrdU). Representative pictures are provided; arrows point at proliferating CMs; scale bars, 30 μm; **(c)** Immunofluorescence analysis of DNA synthesis (BrdU) of post-natal day 7 (P7) cardiomyocytes following stimulation with GR antagonist RU486, alone or in combination with NRG1 (100ng/ml) (n = 6875 cardiomyocytes pooled from the analysis of 18 samples). Cardiomyocytes were identified by cTnI staining and analysed by immunofluorescence for DNA synthesis (BrdU). Representative pictures are provided; arrows point at cardiomyocyte undergoing DNA synthesis; scale bars, 30 μM; (**d**) Immunofluorescence analysis of DNA synthesis (BrdU) of cardiomyocytes derived from adult mice (3 months old – 3M) following stimulation with GR antagonist RU486, alone or in combination with NRG1 (100 ng/ml) (n = 4618 cardiomyocytes pooled from the analysis of 17 samples). Cardiomyocytes were identified by cTnI staining and analysed by immunofluorescence for DNA synthesis (BrdU). Representative pictures are provided; arrows point at cardiomyocytes undergoing DNA synthesis; scale bars 30 μM. The values are presented as mean (error bars show SEM), statistical significance was determined using two-sided Student’s t test in (**a**), (**b**) and using one-way ANOVA followed by Sidak’s test in (**c**), (**d**).

Overall, these findings indicate that GR activity in postmitotic cardiomyocytes suppresses ERK signaling and the mitogenic activity induced by cardiac regenerative growth factors. Pharmacological inhibition of GR restores the ability of these growth factors to effectively activate the MAPK cascade and promote cardiomyocyte proliferation.

### In vivo administration of GR antagonist boosts growth factor-induced cardiomyocyte proliferation and preserves heart function upon anthracycline-induced cardiac damage

Anthracyclines are a cornerstone of many chemotherapy regimens (Curigliano et al., 2016). However, their cumulative and dose-dependent cardiotoxicity limits their use (Morelli et al., 2022) (Cardinale et al., 2015). The mechanisms underlying anthracycline-induced cardiotoxicity are multifaceted, ultimately leading to cardiomyocyte death and loss (Y.-W. Zhang et al., 2009) (S. Zhang et al., 2012) (Linders et al., 2024) (Kim et al., 2018) (Mitry & Edwards, 2016) (Swain et al., 2003). In vitro, doxorubicin, one of the most commonly used anthracyclines, induced cardiomyocyte cell cycle exit in a dose-dependent manner (**Fig. 6a**). Importantly, combinatorial treatment with GR antagonist RU486 and NRG1 significantly increased cardiomyocyte cell-cycle re-entry in neonatal cardiomyocytes pre-treated with doxorubicin (**Fig. 6b**).

**Figure 6.**
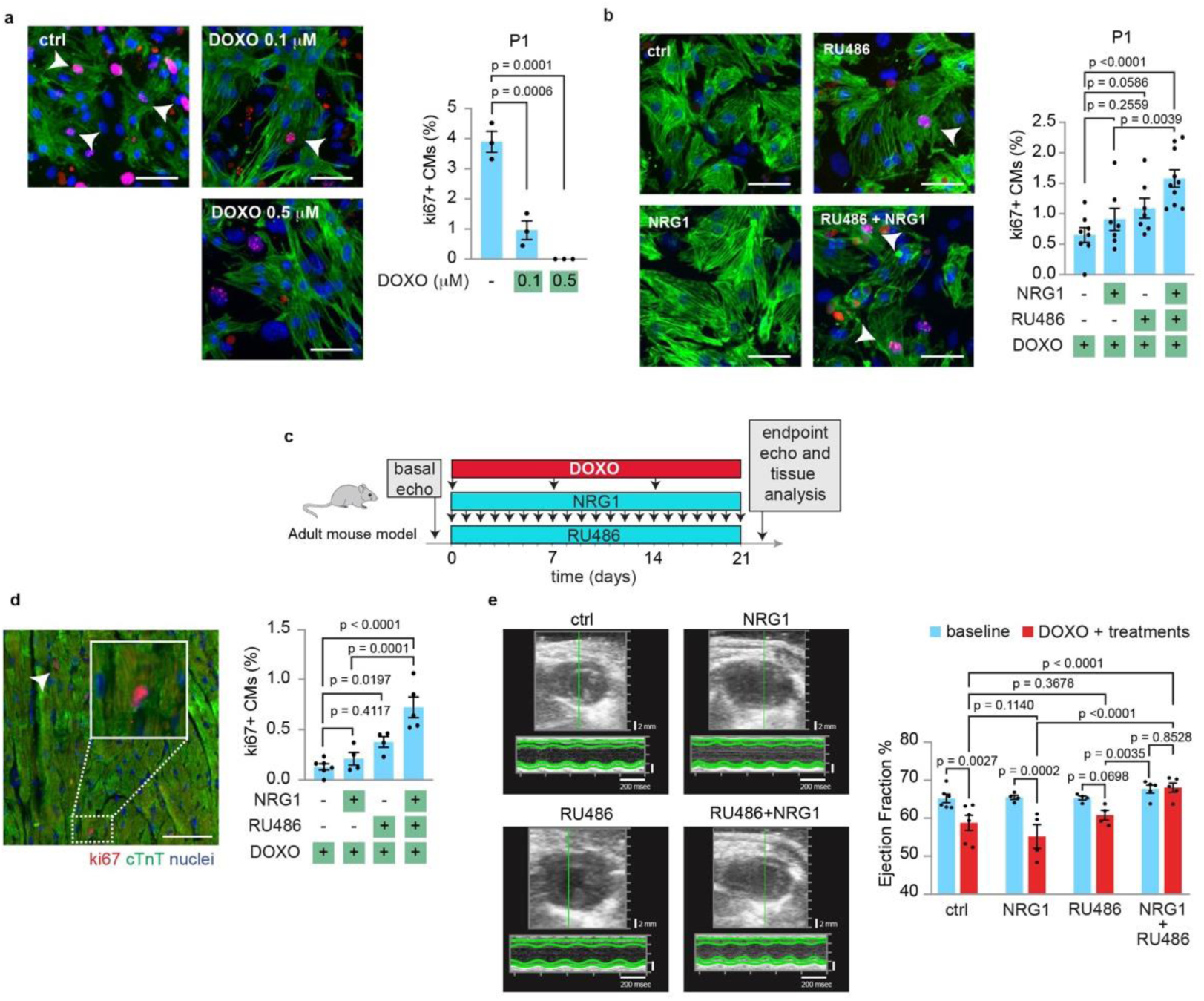
GR antagonization enhances growth factor-induced cardiomyocyte cell-cycle re-entry and heart functional preservation after cardiac damage. (**a**) Immunofluorescence analysis of cell-cycle re-entry (Ki67) of post-natal day 1 (P1) cardiomyocytes following exposure to doxorubicin (DOXO) at different concentrations (0.1 μM and 0.5 μM) (n = 1960 cardiomyocytes pooled from the analysis of 9 samples). Representative figures are provided; white arrows point to cycling cardiomyocytes; scale bar 30 μm; (**b**) Immunofluorescence analysis of cell-cycle re-entry (Ki67) of post-natal day 1 (P1) cardiomyocytes following stimulation with doxorubicin (DOXO, 0.1 μM), alone or in combinations with NRG1 (100ng/ml), RU486 (10^−7^ M), NRG1 (100ng/ml) + RU486 (10^−7^ M) (n = 8725 cardiomyocytes pooled from the analysis of 32 samples). Representative figures are provided; white arrows point cycling cardiomyocytes; scale bar 30 μm; (**c**) Experimental design for the analysis of cardiac damage and regeneration following three IP injections with doxorubicin (DOXO) at 4 mg/kg, once a week (12 mg/kg cumulative dose), in three months old mice. RU486 was provided ad libitum in the feed (20 mg/kg/die) starting after the first injection of DOXO, and NRG1 was administered daily via IP injections (100 ug/kg/die) (control mice received saline). Experimental groups were divided as follows: 1) control group (DOXO alone), 2) DOXO and NRG1 treated, 3) DOXO and RU486 treated, 4) DOXO and RU486+NRG1 treated; (**d)** In vivo evaluation of CM cell cycle re-entry by **i**mmunofluorescence analysis of Ki67 and cardiac Troponin T in heart sections of normal feed (control), NRG1-treated, RU486-treated, RU486+NRG1 treated animals 21 days after doxorubicin (DOXO) (n = 19 mice with a total of 24194 CMs analyzed). Representative pictures of cell cycle activity are provided; arrows point at cycling cardiomyocytes; scale bar, 50 μm; (**e**) Echocardiographic analysis of normal feed (control), NRG1-treated, RU486-treated, RU486+NRG1 treated animals before (basal echo) and after (endpoint echo) 21 days of treatment with a cumulative dose of 12 mg/kg of doxorubicin (DOXO) (n = 19 mice). The values are presented as mean (error bars show SEM), statistical significance was determined using one-way ANOVA followed by Tukey’s test in (**a**), followed by Sidak’s (comparison between pairs of treatments) in (**b**), (**d)** and (**e**).

To test the therapeutic potential of this combination, we took advantage of a mouse model of acute cardiomyopathy induced by doxorubicin. A cumulative dose of 12 mg/kg of doxorubicin was administered via three IP injections, once a week (4 mg/kg on days 0, 7, and 14) (M. Li et al., 2018). RU486 was provided ad libitum through the feed, and NRG1 was administered daily via IP injections (**Fig. 6c**). Analysis of cycling cardiomyocytes by KI67 immunostaining showed that NRG1 alone had minimal effect, while RU486 alone increased cardiomyocyte proliferation (**Fig. 6d**), similarly to the effect we have previously shown in myocardial infarction models (Pianca et al., 2022). Notably, the combination of RU486 and NRG1 significantly boosted cardiomyocyte proliferation (**Fig. 6d**). Echocardiographic assessment 3 weeks after the first doxorubicin administration revealed a significant reduction in ejection fraction in control mice, indicative of impaired cardiac function (**Fig. 6e**). Strikingly, while single-agent treatments showed no significant benefit, the combination of RU486 and NRG1 completely preserved heart function to levels comparable to uninjured mice (**Fig. 6e**). Our results demonstrate that antagonizing endogenous GR activity enhances the efficacy of cardiac regenerative factors *in vivo* following heart injury.

In summary, we here propose a model in which, soon after birth, increased GR activation by glucocorticoids induces *Dusp1* and *Errfi1* expression, which in turn suppress MAPK/ERK signalling and reduce the regenerative potential of growth factors (**Fig. 7**). We suggest transient GR antagonization, for example through RU486 administration, as a promising strategy to enhance the activity of NRG1 and other regenerative growth factors, thereby promoting cardiomyocyte replacement and improving heart function following cardiac injuries (**Fig. 7**).

**Figure 7.**
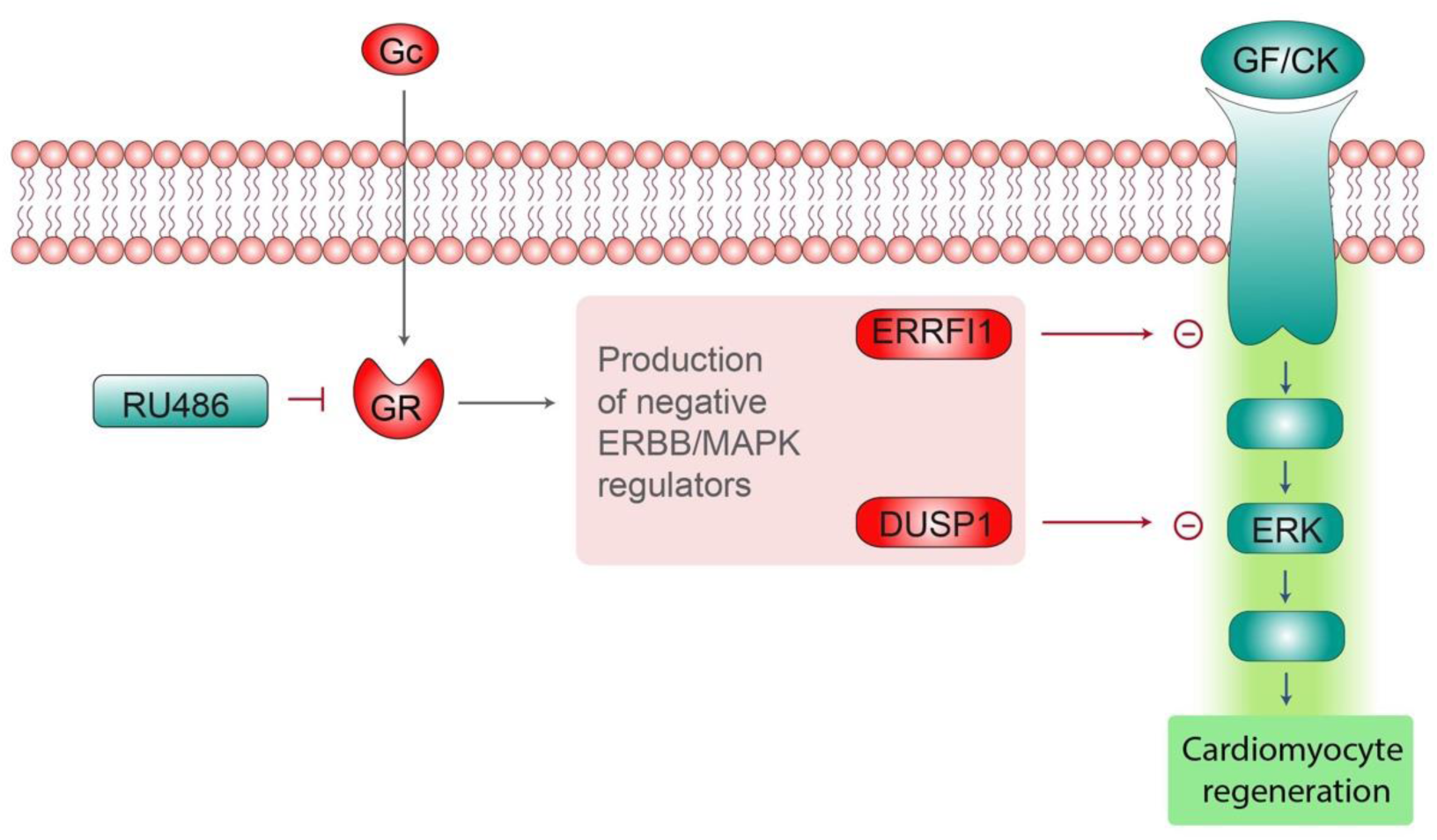
Molecular mechanism of the impact of glucocorticoids on growth factors/cytokines signalling in cardiomyocyte regeneration. Diagram showing the proposed mechanism by which glucocorticoids intercept ERK activation induced by growth factors/cytokines (GF/CK). In detail, glucocorticoids (GCs), upon binding Glucococorticoid Receptor (GR), induce the expression of negative regulators of MAP kinase cascade, namely DUSP1 and ERRFI1. In turn, DUSP1 and ERRFI1 blunt the activation of the MAPK cascade, thereby decreasing growth factors mitogenic potential.

## Discussion

The administration of regenerative growth factors and cytokines is emerging as a promising strategy for heart regeneration. Several cytokines and growth factors suggested for cardiac regeneration signal through the MAPK-ERK pathway (Bongiovanni et al., 2021) (Kubin et al., 2011) (Bongiovanni, Bueno-Levy, et al., 2024) (Wodsedalek et al., 2019) (P. Li et al., 2011). These include receptor tyrosine kinase (RTK)-dependent growth factors such as NRG1 (Bersell et al., 2009) (D’Uva et al., 2015) (Aharonov et al., 2020) (Strash et al., 2021), FGF1 (Engel et al., 2006) (Novoyatleva et al., 2014) (Engel et al., 2005), IGF2 (Shen et al., 2020) (Schuetz et al., 2024b), IGF1 (Sundgren et al., 2003b) (Schuetz et al., 2024a) (Koudstaal et al., 2014), which activate ERK as canonical route, as well as RTK-independent growth factors or cytokines like BMP7 (Bongiovanni, Bueno-Levy, et al., 2024), OSM (Kubin et al., 2011) (Y. Li et al., 2020), LIF (Kubin et al., 2011) (Zou et al., 2003) (Negoro et al., 2001), RANKL (Zacchigna et al., 2018) (Ock et al., 2012), IL6 (Tang et al., 2018) (C. Han et al., 2015b) (Zogbi et al., 2020) (Fahmi et al., 2013), IL13 (Wodsedalek et al., 2019) (Paddock et al., 2021), IL4 (Paddock et al., 2021) (Zogbi et al., 2020) and IL1-β (Palmer et al., 1995) (C. Han et al., 2015b) (Engel et al., 2005) (Przybyt et al., 2013) (Singh et al., 2016), which activate ERK as non-canonical route. In this study, we uncovered a previously unrecognized interplay through which systemic hormones, specifically glucocorticoids, regulate the regenerative potential of cardiac paracrine factors, such as growth factors and cytokines. This effect is mediated by the induction of MAPK/ERK pathway inhibitors, such as DUSP1 and ERRFI1. DUSP1 is a phosphatase that directly inhibits ERK, and its induction likely accounts for the repressive effect of corticosterone on all tested regenerative growth factors and cytokines. In contrast, ERRFI1 has been shown to specifically inhibit ERBB receptors (Anastasi et al., 2016) (Zhong et al., 2021), suggesting that its induction by glucocorticoid action may specifically constrain the NRG1/ERBB2 regenerative axis. Collectively, our findings identify glucocorticoids as critical antagonists of the MAPK/ERK pathway, a pivotal signaling cascade in cardiac regeneration.

Previous studies have shown that ERK signaling dynamics, such as duration, intensity, subcellular localization, feedback mechanisms, and interactions with other pathways, dictate distinct cellular outcomes, including proliferation, survival, and differentiation (Shaul & Seger, 2007) (Von Kriegsheim et al., 2009). Differences in ERK signaling dynamics have been already proven to modulate the regenerative potential across different species and organs (Tomasso et al., 2023) (Yun et al., 2014). Nevertheless, the regulatory mechanisms controlling ERK dynamics in relation to regenerative outcomes remain only partially understood. We propose a model wherein glucocorticoids modulate MAPK/ERK intensity, subcellular localization, and downstream transcriptional responses in cardiomyocytes by inducing DUSP1 and ERRFI1, thereby shifting the outcome from regenerative to non-regenerative. In this regard, DUSP6, a member of the DUSP family, has been shown to attenuate MAPK signaling and limit heart regeneration in zebrafish (Missinato et al., 2018). Moreover, DUSP6, together with other negative feedbacks, has been suggested to contribute to cardiomyocyte redifferentiation, and cell cycle exit after shut-off of ERBB2-induced dedifferentiation program (Shakked et al., 2022). Importantly, our data demonstrate that glucocorticoid-induced upregulation of MAPK/ERK inhibitors and the consequent suppression of growth factor-mediated MAPK/ERK signaling becomes more pronounced during the early postnatal period. This critical developmental window coincides with the transition of cardiomyocytes to a postmitotic state, marked by a significant reduction in their regenerative capacity. During this period, the expression of several pro-regenerative growth factors, such as NRG1, BMP7, IL6, RANKL, and IGF2, declines (Bongiovanni, Bueno-Levy, et al., 2024). Overall, we propose that the diminished mitogenic capacity of endogenous cardiac regenerative factors during the early postnatal period arises from both their reduced production (Bongiovanni, Bueno-Levy, et al., 2024) and the suppression of downstream MAPK signaling (this study). This temporal relationship highlights a developmental context in which glucocorticoids start to actively restrict cardiac regenerative stimuli. Speculatively, this mechanism may drive cardiomyocyte differentiation during early postnatal development and re-differentiation following cardiac injury. However, this hypothesis warrants further investigation.

Our study identifies NRG1 as a model growth factor whose regenerative efficacy is constrained by glucocorticoids. NRG1 administration has shown significant improvements in cardiac function in numerous preclinical studies (Odiete et al., 2012) (De Keulenaer et al., 2019) (X. Liu et al., 2006) (Bersell et al., 2009) (Gu et al., 2010) (Guo et al., 2012) (Bian et al., 2009) (Galindo et al., 2014), establishing its potential as a promising therapeutic strategy for cardiac injury. Modest beneficial effects of NRG1 therapy have also been documented in clinical trials involving heart failure patients (Gao et al., 2010) (Jabbour et al., 2011) (Lenihan et al., 2016). However, these benefits are most likely independent of cardiomyocyte proliferation, as NRG1 ability to stimulate cardiomyocyte proliferation is predominantly restricted to the neonatal stage (D’Uva et al., 2015) (Polizzotti et al., 2015) (Reuter et al., 2014). To date, the limited mitogenic potential of NRG1 in adult mammals has been primarily attributed to the low expression of its receptor, ERBB2, in adult cardiomyocytes (D’Uva et al., 2015). Our findings provide new insights into this limitation, revealing an additional regulatory mechanism: the glucocorticoid-dependent upregulation of MAPK inhibitors such as DUSP1 and ERRFI1. We propose that transient antagonism of GR activity could amplify the regenerative efficacy of NRG1. In support of this, combining RU486, a GR antagonist, with NRG1 in a mouse model of doxorubicin-induced cardiomyopathy resulted in a synergistic enhancement of cardiomyocyte proliferation and functional preservation, markedly surpassing the effects observed with either treatment alone. This combinatorial approach underscores the potential of transient GR antagonism to improve the efficacy of growth factor-based regenerative therapies. Beyond NRG1, we demonstrate that glucocorticoids similarly inhibit the regenerative capacity of several other growth factors, including BMP7, IL6, IGF2, and FGF1, further broadening the scope of this strategy.

Our findings also have implications for other regenerative contexts. Indeed, the MAPK/ERK pathway is emerging as a universal regenerative hub across multiple organs and species (De Simone et al., 2021) (Tomasso et al., 2023) (Wen et al., 2022) (Xiao & Xiong, 2023) (X.-S. Zhang et al., 2024) (Yun et al., 2014), with roles in the regeneration of the heart (P. Han et al., 2014) (Missinato et al., 2018) (P. Liu & Zhong, 2017), liver (Ohashi et al., 2021), bone (De Simone et al., 2021), optic nerve (Duprey-Díaz et al., 2016) (Yasumuro et al., 2017), retina (Duprey-Díaz et al., 2016) (Yasumuro et al., 2017), limbs (Yun et al., 2014) (Suzuki et al., 2007) (Blassberg et al., 2011), and fin (X.-S. Zhang et al., 2024) among others. This pathway even facilitates whole-body regeneration in invertebrates (Fan et al., 2023). Interestingly, glucocorticoids have been implicated as inhibitors of regeneration in several non-cardiac tissues, including bone, hair follicle and nervous system (Hachemi et al., 2018) (Choi et al., 2021) (Kyritsis et al., 2012). These parallels suggest that the interplay between glucocorticoids and MAPK/ERK signaling may represent a conserved mechanism that limits regeneration across multiple organ systems. These findings are consistent with our previous work in breast cells, where glucocorticoids suppressed EGFR signaling by upregulating negative feedback regulators DUSP1 and ERRFI1 (Lauriola et al., 2014). Together, these studies highlight a broader paradigm in which glucocorticoids fine-tune growth factor signaling by reinforcing negative feedback mechanisms.

In conclusion, our study provides a mechanistic framework to explain how systemic glucocorticoids suppress cardiac regeneration and highlights the therapeutic potential of transient GR antagonism. By enhancing the regenerative capacity of growth factor-based therapies, this approach offers a promising strategy for addressing the unmet need in heart failure treatment.

## Materials and methods

### Mouse neonatal and juvenile cardiomyocyte isolation and culture

Primary neonatal cardiomyocytes were extracted from the hearts of 0-day-old (P0) or 1-day-old (P1) mice. Neonatal cardiac cells were isolated by enzymatic digestion with pancreatin (Sigma) and collagenase (Roche), as we recently described (Bongiovanni, Miano, et al., 2024). Primary juvenile cardiomyocytes were extracted from 7-day-old (P7) mouse hearts which were anterogradely perfused with digestion buffer containing collagenase (Roche), trypsin (Sigma), and protease (Sigma), as previously described (Omatsu-Kanbe et al., 2018). Cells were then cultured in 0.1% gelatine-coated (Sigma) wells with DMEM/F12 (Aurogene) supplemented with 1% L-glutamine (Sigma), 1% sodium pyruvate (Life Technologies), 1% non-essential amino acids (Life Technologies), 1% penicillin and streptomycin (Euroclone), 5% horse serum (Invitrogen) and 10% FBS (Life Technologies) (hereafter referred to as ‘complete-medium’) at 37°C and 5% CO2. The cells were allowed to adhere for 48 h in complete-medium (24 h for knockdown analysis). Subsequently, the medium was replaced with an FBS-deprived complete-medium containing selected growth factors (NRG1, FGF1, IGF2, IGF1, BMP7, OSM, LIF, RANKL, IL6, IL13, IL4, IL1-beta, Immunotools), corticosterone (27840, Sigma), mifepristone (RU486, Sigma, M8046) or BCI (317496, Sigma) for about 48 h (for proliferation analyses) or at specific time points for cell signaling analysis. For BrdU assays, BrdU (10 mM, B5002, Sigma) was introduced along with the treatments. For gene or protein expression analysis, 24h of starvation in serum-deprived medium was performed before cell treatment.

### Mouse adult cardiomyocyte isolation and culture

Primary adult cardiomyocytes were obtained from 3-4 months old mice as previously described (Ackers-Johnson et al., 2016). Following cervical dislocation, EDTA buffer was injected into the right ventricle to prevent clotting. The ascending aorta was clamped and the heart excised. The heart was then anterogradely perfused with EDTA buffer, perfusion buffer, and collagenase buffer through the left ventricle to digest the tissue. Once digestion was complete, the clamp was removed, and the tissue was gently broken into pieces of roughly 1 mm x 1 mm, followed by gentle trituration for 2 minutes using a serological pipette. Enzymatic digestion was stopped and the cell suspension was filtered through a 100 μm strainer. Gradual calcium reintroduction was then achieved through sequential steps of gravity settling and resuspension in perfusion buffer mixed with increasing concentrations of culture medium. For each heart, the final myocyte pellet was resuspended in 2 ml of room temperature culture medium and counted. Cells were then plated in 10 ug/ml fibronectin-coated (Sigma) wells in M199 (Sigma) supplemented with 5% FBS, 10 mM BDM (Sigma) and 1% penicillin and streptomycin (Euroclone) (hereafter referred to as ‘plating medium’) at 37°C and 5% CO2. After 1 h, the plating medium was removed and cells were incubated for 48h in M199 supplemented with 10 mM BDM, 1% penicillin and streptomycin, 0.1% BSA, 1% ITS (Sigma) and 1% chemically defined lipid concentrate (Sigma) (hereafter referred to as ‘culture medium’). Subsequently, the culture medium was replaced with fresh one supplemented with NRG1 or mifepristone for about 48 h. For BrdU assays, BrdU was introduced along with the treatments.

### Cardiomyocyte and stromal cell separation

Cardiomyocytes were separated from cardiac stromal cells using the MACS Neonatal Isolation System (Miltenyi Biotech#130-100-825), following the manufacturer’s protocol. The isolation kit includes a cocktail of monoclonal antibodies conjugated with MACS® MicroBeads, designed to magnetically retain non-cardiomyocyte cells within a MACS Column placed in the magnetic field of a MACS Separator. Enriched cardiomyocytes were then cultured in a complete medium. The validation of the enrichment procedure was previously demonstrated by the analysis of gene expression levels of markers specific for cardiomyocytes and stromal cells (fibroblasts and endothelial cells) (Pianca et al., 2022).

### Transient gene knockdown

To silence specific molecular targets, SMARTpool siRNAs targeting Dusp1 (Dharmacon#L-040753-00-0005) and Errfi1 (Dharmacon#L-040753-00-0005) were delivered to postnatal day 1 (P1) cardiomyocytes, 24 hours after seeding, using Lipofectamine™ 3000 Reagent (Thermo Fisher #L3000008), following the manufacturer’s protocol. Lipofectamine 3000 Reagent was used at the lowest recommended concentration. Gene knockdown was obtained following 48h of transfection at 37°C and 5% CO2. Subsequently, cells were lysed for RNA or protein extraction, as described below, or the medium was replaced with an FBS-deprived complete-medium containing NRG1 and corticosterone for 48 h (for proliferation assays).

### Immunofluorescence on primary cultured cardiomyocytes and cardiac tissue sections

Cultured cells were fixed with 4% paraformaldehyde (PFA) solution (Sigma, diluted in PBS) for 20 min at 4°C and then washed three times with PBS. Hearts were fixed in 4% PFA for 24h, dehydrated and embedded in paraffin. 4 µm-thick sections were deparaffinized, rehydrated and boiled in EDTA buffer at pH 8.5 for antigen retrieval. Both tissue sections and cultured cells were then permeabilized with 0.5% Triton-X100 (Sigma) in PBS for 5 minutes at room temperature. Non-specific antibody binding was prevented by applying a blocking solution of PBS supplemented with 5% BSA or 5% goat serum with 0.1% Triton-X100 for 1 hour at room temperature. The samples were subsequently incubated overnight at 4°C in a humidified chamber with primary antibodies diluted in PBS, supplemented with 3% BSA or 3% goat serum and 0.1% Triton-X100. The following primary antibodies were used: anti-cardiac Troponin T (1:800, ab33589, abcam), anti-cardiac Troponin I antibody (1:800, ab47003, abcam), anti-Ki67 (1:50, ab16667, abcam), anti-BrdU (1:40, G3G4, DSHB), anti-ERK2 (1:50, sc-1647, Santa Cruz), anti-phospho-ERK (1:400, M8159, Sigma) and anti-AIM1 (1:50, 611082, BD Biosciences). After primary antibody incubation, the samples were washed three times with PBS and then incubated for 1 hour at room temperature with fluorophore-conjugated secondary antibodies (Alexa Fluor 488-AffiniPure Goat-Anti Mouse/Rabbit IgG or Cy3-AffiniPure Goat-Anti-Mouse/Rabbit IgG, Jackson ImmunoResearch Labs) diluted in PBS supplemented with 1% BSA or 1% goat serum, and 0.1% Triton-X100. After three washes in PBS, DAPI (4’,6-diamidino-2-phenylindole dihydrochloride, Sigma), diluted to 1 µg/ml in PBS, was applied for 10 minutes at room temperature to visualize the nuclei. The slides were then mounted with an antifade solution (Vectorlabs), covered with a coverslip, sealed with nail polish, and imaged. Slides were imaged at an Olympus VS200 slide scanner at 20x magnification.

### Protein Extraction and Western Blotting

400,000 cells were seed for protein analysis experiments. Cells were lysed with RIPA buffer supplemented with protease inhibitor (Sigma) and phosphatase inhibitors (Sigma). Then, 25 µg of protein extracts were resolved by sodium dodecyl sulfate (SDS)-polyacrylamide gel electrophoresis and transferred to a nitrocellulose membrane (AmershamTM ProtranTM Premium, 0.45 mm). The membrane was blocked for 1 h using TBS-T (0.1% Tween 20) supplemented with 5% BSA (Sigma), and incubated overnight at 4°C with the following primary antibodies: anti-phospho-GR (Ser211) (1:1000, 4161, Cell Signaling), anti-GR (1:1000, 12041, Cell Signaling), anti-DUSP1 (1:250, sc-373841, Santa Cruz), anti-ERRFI1 (1:250, 137152, Santa Cruz), anti-ERK2 (1:1000, sc-1647, Santa Cruz), anti-phospho-ERK (1:10000, M8159, Sigma), anti-AKT (1:1000, 9272, Cell Signaling), anti-phospho-AKT (1:1000, 9271, Cell Signaling), anti-GAPDH (1:1000, 2118, Cell Signaling) and anti-b-actin (1:500, sc-4778, Santa Cruz). For protein detection, the membrane was incubated with anti-rabbit or anti-mouse horseradish peroxidase-conjugated secondary antibody (Dako EnVision+ System-HRP Labeled Polymer) followed by a chemiluminescent reaction (Clarity Western ECL Substrate, Bio-Rad). Signals and images were acquired by ChemiDoc XRS 2015 (Bio-Rad Laboratories), and densitometric analysis were performed using Image Lab software (version 5.2.1; Bio-Rad Laboratories).

### RNA sequencing

Transcriptomic analysis was performed using QuantSeq FWD 3′ mRNA-Seq Services of Lexogen. In brief, purified total RNA was extracted from in vitro cultures of cardiomyocytes and from whole heart homogenates; single-read sequencing was performed on a NextSeq 500 (Illumina), and quantification was based on tags in the 3′ region with more than 20 million reads per sample. Data analysis was performed with the following software packages: NextSeq Control Software version 2.2.0.4 (for base-calling), BCL-to-FASTQ file converter bcl2fastq (for demultiplexing), cutadapt version 1.16 (for trimming), FastQC version 0.11.7, multiqc version 1.5, STAR version 2.5.4a (for alignment), featureCounts version 1.6.2 (for counting) and DESeq2 version 1.18.1 (for differential expression). Gene Ontology (GO) analysis was conducted using the PANTHER Classification System (Release 17.0), available at http://www.pantherdb.org/ (Mi et al., 2019). A statistical overrepresentation test was applied to identify significantly enriched GO terms within the GO-Biological Process annotation set. Significantly upregulated genes (adj. p-value < 0.05) from RNA-seq data of corticosterone-treated samples were used as input. The heat maps shown in Fig. 2 and Fig. 4 were generated using the heatmap2 Galaxy package.

### Transcriptional analysis

Total RNA extraction was performed with the NucleoSpin RNA II kit (Macherey Nagel) according to the manufacturer’s protocol. RNA quantification and quality check were performed using a Nanodrop spectrophotometer (N1000, Thermo). RNA was reverse transcribed to double-stranded cDNA using the RevertAid RT Kit (Thermo) according to the manufacturer’s protocol. Real-Time (rt)-PCR was performed using Fast SYBR Green PCR Master Mix (Applied Biosystems) on a QuantStudio 5 Flex instrument (Applied Biosystems). Oligonucleotide sequences of genes analyzed in this study are listed in the table below. Relative quantification was performed using the Hprt1 gene as a loading control. DDCT was calculated and data of each gene were analyzed using a 2DDCT method and reported as mean fold change.

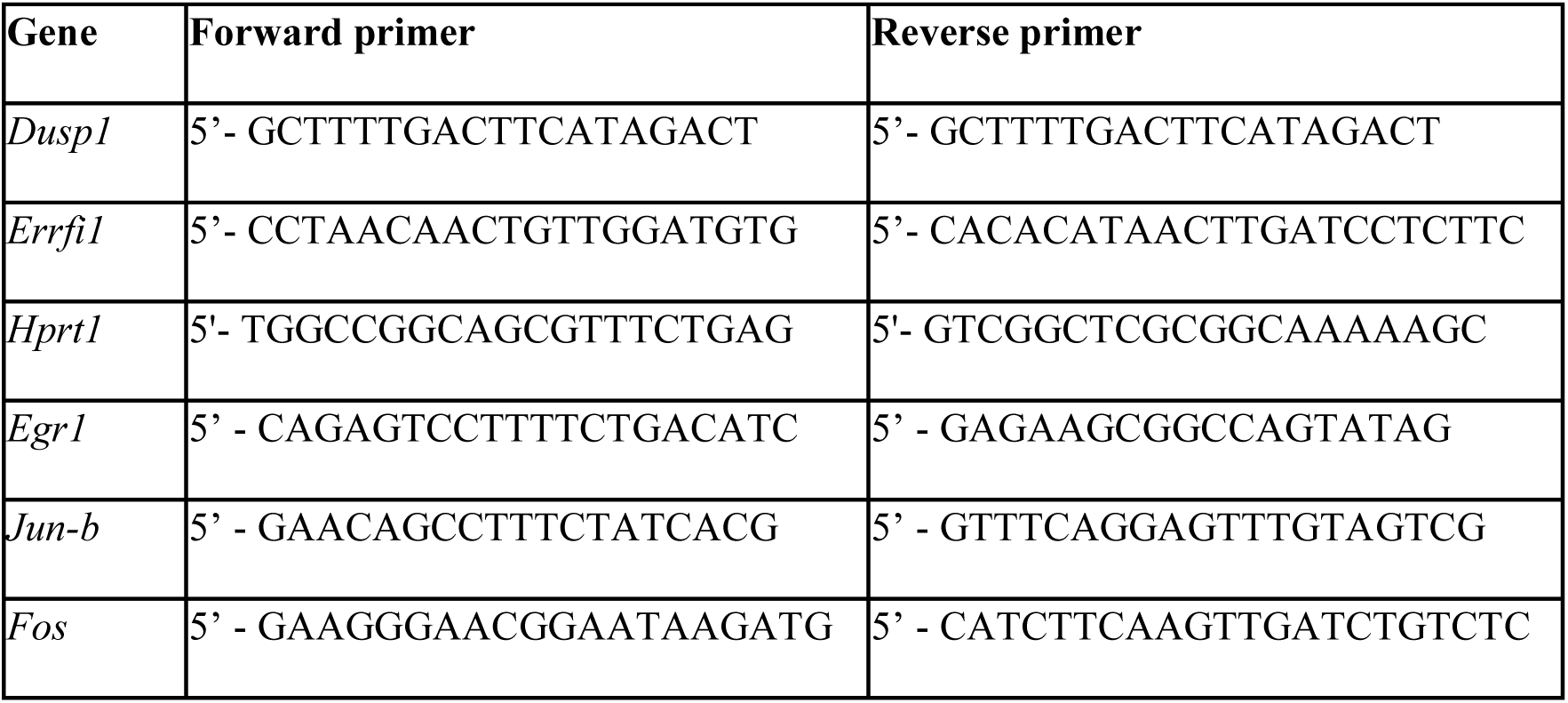

### Time-lapse imaging

Following cell isolation, postnatal day 1 (P1) cardiomyocytes were seeded at 35,000 cells/well in a 96-well plate and left to adhere for 48 h. Then, the complete medium was replaced with serum-free medium supplemented with 10 nM TMRE (tetramethylrhodamine ethyl ester, Sigma), a cationic fluorescent dye that labels active mitochondria and therefore allows to selectively visualize cardiomyocytes (Hattori et al., 2010). After 20 min of incubation at 37°C, the medium containing TMRE was removed, and cardiomyocytes were treated with NRG1 (100 ng/mL, R&D) and corticosterone (10^−8^ M) in FBS-deprived complete medium. Live cell imaging was performed using a widefield fluorescent microscope (Nikon Eclipse Ti2). Timelapse imaging was carried out for about 16 h and images were acquired at x40 magnification every 20 min.

### Mouse experiments

Mice were housed at a controlled temperature of 20–25 °C and a relative humidity of 40–60%, following a 12-hour light/dark cycle. Experiments were approved by the Animal Care and Use Committee of the University of Bologna and Weizmann Institute of Science. Mice studies were performed on C57BL/6 background, cardiomyocyte-specific GR knock-out (GR-cKO) and cardiomyocyte-specific costitutively active ERBB2 (caERBB2) mice. Cardiomyocyte-specific GR knock-out (GR-cKO) mice were generated as previously reported (Pianca et al., 2022).

Cardiomyocyte-restricted Dox-inducible overexpression of ca*Erbb2* was generated as before (D’Uva et al., 2015). Doxycycline (TD02503, Harlan Laboratories) was administered in the food to repress transgene expression. Mice’s age is reported in the main text, figures and/or figure legends. Data were aggregated without discriminating for gender.

### Doxorubicin-induced myocardial injury

The experimental setups are depicted in Fig. 6. A total of nineteen C57BL/6J mice of indicated genotypes were divided into four groups: doxorubicin alone, NRG1 + doxorubicin, RU486 + doxorubicin, NRG1 + RU486 + doxorubicin. All nineteen mice received intraperitoneal (i.p.) injections of doxorubicin (ab120629, abcam) at a cumulative dose of 12 mg/kg, administered via 3 injections, once a week (4 mg/kg on days 0, 7, and 14) (M. Li et al., 2018).

### Echocardiography

Heart function was evaluated in adult mice before and after doxorubicin-induced myocardial injury by transthoracic echocardiography performed on sedated mice (isoflurane 1.5%) using a MyLab 70 XV device. Parasternal Short Axis view was used to obtain measures to calculate ejection fraction (EF) and fractional shortening (FS).

### In vivo drug delivery

In the experiment analyzing the effects of GR blockage during physiologic postnatal development, GR antagonist mifepristone (RU486, Sigma-Aldrich, M8046) was dissolved in the drinking water (42 μg/ml, equivalent to 20 mg/kg/day). Freshly prepared solution was made every day and administered ad libitum to the lactating mother, starting from the day of the birth of the pups and continuing the treatment for 7 days, as previously done (Pianca et al., 2022). Based on previous studies (Su et al., 2015), the amount of RU486 received by pups is expected to be around 0.3 mg/kg.

In the experiment analyzing the effects of GR blockage and NRG1 after doxorubicin-induced cardiac damage in adulthood, GR antagonist mifepristone (RU486, Sigma-Aldrich, M8046) was delivered through the food (special food diet dry pellets were produced by a commercial manufacturer, Mucedola), with a dose of 20 mg/kg/day, starting the administration immediately after the first injection of doxorubicin. NRG1 (Biotechne, 396-HB) was administered via daily IP injections at 100 ug/kg. Meanwhile, other groups were i.p. injected with an equal volume of saline solution.

### Statistical analyses

Analyses were conducted using GraphPad Prism 8 software. Whenever normality could be assumed, the two-sided Student’s t-test or analysis of variance (ANOVA) followed by Sidak’s, Tukey’s and Dunnet’s test were used to compare group means, as specified in the figure legends. A P value < 0.05 was considered statistically significant. In all panels, numerical data are presented as the mean ± standard error of the mean (s.e.m.). Every dot represents a different biological replicate, as indicated in the figure legends, together with the number of cardiac cells analyzed.

### Data availability

The datasets generated and/or analyzed in this study are available in the source data file. RNA sequencing data were deposited in the Gene Expression Omnibus repository under accession numbers GSE286562 (controls in GSE202968).

## Acknowledgments

We acknowledge financial support under the National Recovery and Resilience Plan (NRRP), Mission 4, Component 2, Investment 1.1, Call for tender No. 1409 published on 14.9.2022 by the Italian Ministry of University and Research (MUR), funded by the European Union – NextGenerationEU– Project Title “The cardiomyocyte-intrinsic role of the glucocorticoid receptor in cardiac aging” – CUP J53D23018140001 Grant Assignment Decree No. 1369 adopted on 01/09/2023 by the Italian Ministry of Ministry of University and Research (MUR).

The research was also supported by the European Union’s Horizon 2020 research and innovation programme under the ERA-NET on Cardiovascular Diseases (ERA-CVD) Co-fund action to G.D’U. and E.T. (Grant Number: JCT2016-40-080), Fondazione Carisbo to G.D’U. (Grant number: 2023.0210) and by the Italian Ministry of Health (RC-2024-2790614). The views and opinions expressed are those of the authors only and do not necessarily reflect those of the European Union or the European Commission. Neither the European Union nor the European Commission can be responsible for them.

S.D.P. was supported by a PON Research and Innovation 2014–2020 (FSE React-EU) PhD Scholarship (code DOT1303972 - CUP J35F21003360006) funded by the Italian Ministry of University and Research (MUR) under DM 1061/2021, Action IV.4 “Doctorates and Research Contracts on Innovation Topics”.

E.T. was supported by the European Research Council (ERC AdG #788194).

