## Supplementary Figures and Tables for "Harnessing glucocorticoid receptor antagonization to enhance the efficacy of cardiac regenerative growth factors and cytokines"

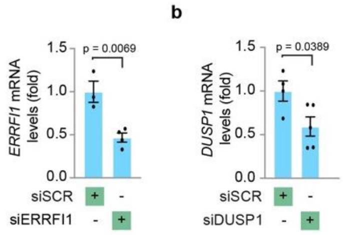


**Supplementary Figure 1.** ***In vitro*** **analysis of the efficacy of** ***Errfi1 and Dusp1*** **gene knockdown in neonatal cardiomyocyte cultures. (a-b)** mRNA expression levels of **(a)** *Errfi1* (n = 3 for siSCR; n = 4 for siERRFI1) and **(b)** *Dusp1* (n = 4 for siSCR; n = 5 for siDUSP1) from cultured postnatal day 1 (P1) cardiomyocytes following siRNA delivery for 48 hours. In all panels, numerical data are presented as mean (error bars show s.e.m.). Statistical significance was determined using two-sided Student's t-test in (**a-b**).

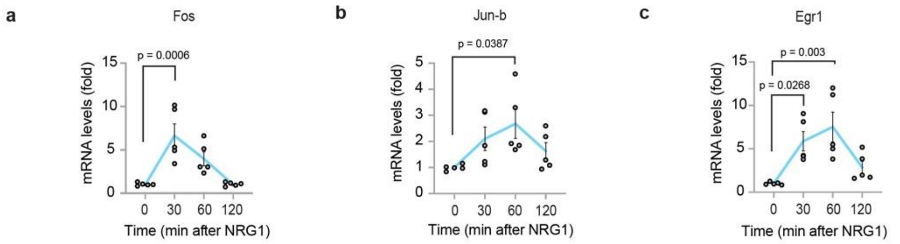


**Supplementary Figure 2. Activation of NRG1 signalling leads to the expression of IEGs in cardiomyocytes.** (**a-c**) Real-time PCR analysis of IEGs (Immediate Early Genes), namely (**a**) *Fos,* (**b**) *Jun-b,* and (**c**) *Egr1*, upon treatment with NRG1 (100 ng/ml) for 30, 60 and 120 minutes (n = 5 replicates per time point) In all panels, numerical data are presented as mean (error bars show s.e.m.). Statistical significance was determined by one way ANOVA followed by Tukey's test in (**a-c**).

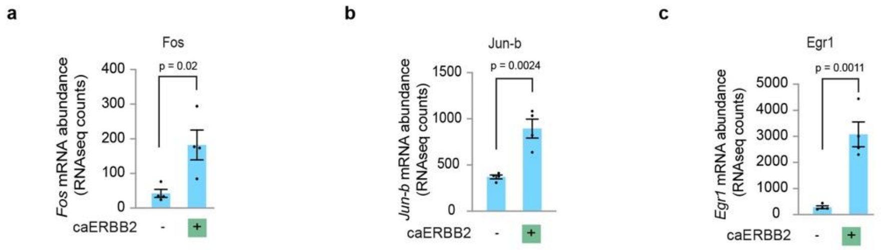


**Supplementary Figure 3. Activation of ERBB2 signalling in postmitotic cardiomyocytes leads to the expression of IEGs. (a-c)** Analysis of (**a**) Fos, (**b**) Jun-b and (**c**) Egr1 expression from RNA-sequencing data of cardiac tissue of a mouse model with cardiomyocyte-restricted overexpression of a constitutively active ERBB2 (caERBB2) isoform (Aharonov et al., 2020) (n = 4 replicates wild type adult mice; n = 4 replicates caERBB2 adult mice). In all panels, numerical data are presented as mean (error bars show s.e.m.). Statistical significance was determined by two-sided Student's t-test in (**a-c**).

**Supplementary Table 1. List GR target genes in cardiomyocytes.** List of significantly upregulated genes (Adj pvalue < 0.05, LogFC > 0) by RNA-Seq analysis of cultured neonatal cardiomyocytes treated *in vitro* with corticosterone (CORT 10^-6M).

| Ensembl ID | Gene symbol | Log2 fold change | Adjusted p value |
|---|---|---|---|
| ENSMUSG00000020893 | Per1 | 1.875938 | 1.59E-16 |
| ENSMUSG00000048489 | 8430408G22Rik | 1.447621 | 1.37E-13 |
| ENSMUSG00000022893 | Adamts1 | 0.843179 | 1.37E-13 |
| ENSMUSG00000021025 | Nfkbia | 0.833226 | 1.37E-13 |
| ENSMUSG00000019970 | Sgk1 | 0.891416 | 1.23E-10 |
| ENSMUSG00000048546 | Tob2 | 1.109562 | 3.53E-10 |
| ENSMUSG00000035828 | Pim3 | 0.703874 | 1.41E-09 |
| ENSMUSG00000071637 | Cebpd | 0.738387 | 1.89E-09 |
| ENSMUSG00000024190 | Dusp1 | 0.9453 | 1.52E-08 |
| ENSMUSG00000020108 | Ddit4 | 0.775985 | 8.48E-08 |
| ENSMUSG00000033863 | Klf9 | 0.619398 | 1.06E-07 |
| ENSMUSG00000032515 | Csrnp1 | 0.877104 | 6.45E-07 |
| ENSMUSG00000000078 | Klf6 | 0.459213 | 2.90E-06 |
| ENSMUSG00000031765 | Mt1 | 0.464555 | 1.16E-05 |
| ENSMUSG00000028967 | Errfi1 | 0.930957 | 4.06E-05 |
| ENSMUSG00000031431 | Tsc22d3 | 0.474027 | 0.001177324 |
| ENSMUSG00000032712 | 2810474O19Rik | 0.500258 | 0.001568283 |
| ENSMUSG00000002910 | Arrdc2 | 1.710514 | 0.001602811 |
| ENSMUSG00000046694 | Fam46b | 0.803375 | 0.002193925 |
| ENSMUSG00000026360 | Rgs2 | 0.600711 | 0.002481857 |
| ENSMUSG00000021268 | Meg3 | 0.248672 | 0.003362267 |
| ENSMUSG00000092274 | Neat1 | 0.318677 | 0.003549614 |
| ENSMUSG00000025372 | Baiap2 | 0.414556 | 0.006887199 |
| ENSMUSG00000061878 | Sphk1 | 1.110568 | 0.00731603 |
| ENSMUSG00000031762 | Mt2 | 0.910539 | 0.007490008 |
| ENSMUSG00000101603 | Gm28730 | 0.664942 | 0.008328156 |
| ENSMUSG00000105431 | Gm42640 | 1.666903 | 0.009579226 |
| ENSMUSG00000065503 | Mir351 | 0.736576 | 0.011042489 |
| ENSMUSG00000024758 | Rtn3 | 0.232608 | 0.018194425 |
| ENSMUSG00000073684 | Faap20 | 0.391626 | 0.020384986 |
| ENSMUSG00000030867 | Plk1 | 0.297872 | 0.02519733 |
| ENSMUSG00000032578 | Cish | 0.418896 | 0.026079025 |
| ENSMUSG00000021453 | Gadd45g | 0.295872 | 0.026352559 |
| ENSMUSG00000103469 | Gm9910 | 4.76356 | 0.034968173 |
| ENSMUSG00000026435 | Slc45a3 | 1.567113 | 0.034968173 |
| ENSMUSG00000037411 | Serpine1 | 0.242284 | 0.034968173 |

| | | | |
|---|---|---|---|
| ENSMUSG00000055148 | Klf2 | 0.33775 | 0.035416601 |
| ENSMUSG00000114501 | AC154176.2 | 0.986415 | 0.04497072 |
| ENSMUSG00000030790 | Adm | 0.325775 | 0.04497072 |
| ENSMUSG00000032300 | 1700017B05Rik | 0.483477 | 0.048242628 |
